# Maize root growth, Oxygen and N availability drive formation of N_2_O hotspots in soil

**DOI:** 10.1101/2025.06.03.657623

**Authors:** Pauline Sophie Rummel, Martin Reinhard Rasmussen, Aurélien Saghaï, Theresa Merl, Sara Hallin, Carsten W. Mueller, Klaus Koren

## Abstract

Plant roots modify all major controls of denitrification in soils, particularly the availability of the main substrates (NO_3_^−^ and C_org_), soil moisture, soil O_2_ content, and root-associated microbial communities, and thus play an important role in N_2_O formation. Direct in-situ measurements of N_2_O concentrations in the rhizosphere are lacking, yet crucial to understanding how rhizosphere denitrification contributes to overall N_2_O emissions from soil. We equipped rhizoboxes with O_2_-sensitive planar optodes to simultaneously monitor root growth and rhizosphere/soil O_2_ concentrations. We measured soil surface N_2_O fluxes and linked them to root growth, soil moisture, and root/soil O_2_ concentrations. Based on root growth and O_2_ concentrations, we identified regions of interest (ROI) and sampled small soil volumes, which were analyzed for C, N, abundance of microbial denitrifiers (*nirK, nirS*) and N_2_O reducers (*nosZI, nosZII*), and soil N_2_O concentrations. Plant roots determined depth gradients of nutrients and denitrification gene abundances in the soil of the rhizoboxes with higher resource availability (NO_3_^-^, DOC) and lower soil moisture in the upper soil layers, which also had higher abundances of total bacteria, *nirK* and *nosZII*. We anticipate that these uppermost soil layers largely contributed to N_2_O formation. For the first time we were able to show high *in-situ* N_2_O concentrations with distinct depth profiles around roots, and O_2_ and N availability controlling N_2_O production at the process scale.

## Introduction

Plant roots play an important role in regulating denitrification, the main process contributing to formation of the climate-relevant greenhouse gas nitrous oxide (N_2_O) (Butterbach-Bahl *et al*., 2013; Ciais *et al*., 2013). Denitrification is the sequential reduction of nitrate (NO_3_^-^) to nitrite (NO_2_^-^), nitric oxide (NO), nitrous oxide (N_2_O), and molecular nitrogen (N_2_) by heterotrophic bacteria, fungi, and archaea (Zumft, 1997). Plant roots modify major controls of denitrification through altering soil moisture, O_2_ content, the availability of its main substrates (NO_3_^−^ and C_org_), and root-associated microbial communities. While the rhizosphere is indicated to be a hotspot of denitrification, the extent of N_2_O formation in the rhizosphere and its contribution to N_2_O emissions from soils remain unclear.

Increased N_2_O losses have been reported from planted compared to bare soils (Woldendorp, 1963; Vinther, 1984; Senbayram *et al*., 2020; Yankelzon *et al*., 2024b) and the promoting effect of plant growth on denitrification has been associated with higher C_org_ availability in the rhizosphere (Smith & Tiedje, 1979; Bakken, 1988; Philippot *et al*., 2009). Plants translocate 20-30 % of assimilated C belowground and about 7-11 % of assimilated C enters the soil as rhizodeposition (Pausch & Kuzyakov, 2018). Higher C availability in the rhizosphere attracts soil microorganisms including bacteria, fungi, oomycetes, and archaea feeding on rhizodeposits making the rhizosphere a hotspot of microbial activity (Starkey, 1958; Philippot *et al*., 2013). In addition to microbial respiration, root respiration largely contributes to O_2_ consumption in soil with rates of around 20 mg O_2_ m^−1^ root d^−1^ for maize roots (Ben-Noah & Friedman, 2018). Root and microbial respiration decrease O_2_ availability, creating suitable conditions for denitrification (Bateman & Baggs, 2005; Hu *et al*., 2015). Accordingly, abundance and composition of bacterial communities differ significantly between planted and unplanted soil, and between bulk soil and the rhizosphere (Chèneby *et al*., 2004; Henry *et al*., 2008; Philippot *et al*., 2013). Studies show that denitrifiers are more abundant in the rhizosphere than the bulk soil of a range of different plant species (Knowles, 1982; Chèneby *et al*., 2004; Ai *et al*., 2020; Wang *et al*., 2024). Accordingly, nitrate reduction and denitrification activities were higher in rhizosphere soil compared to bulk soil (Smith & Tiedje, 1979; Klemedtsson *et al*., 1987; Bakken, 1988; Mahmood *et al*., 1997; Hamonts *et al*., 2013; Guyonnet *et al*., 2017; Malique *et al*., 2019; Zhao *et al*., 2020).

Direct in-situ measurements of N_2_O concentrations in the rhizosphere are lacking, yet crucial to understanding how roots control denitrification and N_2_O formation. Further, it remains unknown to which extent rhizosphere denitrification contributes to overall N_2_O emissions from soil and how microscale N_2_O production relates to surface N_2_O fluxes. Denitrification takes place in microbial hotspots – small soil volumes with faster process rates compared to the average soil conditions (=coldspots) (Groffman *et al*., 2009; Kuzyakov & Blagodatskaya, 2015). Characterization of these N_2_O hot and cold spots is needed to better understand the ‘rhizosphere effect’ and its significance for N_2_O emissions.

This study aimed (I) to elucidate how plant roots control formation of N_2_O hotspots in the rhizosphere, (II) to determine spatial distribution of N_2_O production (hotspots) in relation to root growth and development, and (III) to characterize microbial N cycling in N_2_O hotspots. We hypothesize that (I) root growth promotes microbial growth and respiration, increasing O_2_ consumption at roots and in the rhizosphere. Accordingly, N_2_O fluxes from soil increase with increasing root growth. (II) Increased microbial respiration from root exudation stimulates formation of anoxic microsites (N_2_O hotspots) in close proximity of the roots. These N_2_O hotspots are characterized by higher availability of C and N, and higher abundance of N reducing microorganisms compared to cold spots with low N_2_O concentrations. To address these hypotheses, we equipped rhizoboxes with O_2_-sensitive planar optodes to simultaneously monitor root growth and rhizosphere/soil O_2_ concentrations. We further measured surface N_2_O fluxes and linked them to root growth, soil moisture, and root/soil O_2_ concentrations. Finaly, we defined regions of interest (ROI) related to root growth and O_2_ concentrations, and analyzed them for C, N, abundance of microbial N reducers, and soil N_2_O concentrations.

## Material and Methods

### Optode fabrication

Planar optodes are a non-invasive imaging technique based on reversible changes in luminescence properties of analyte-specific fluorophores enabling visualization and quantification of biochemical processes with an emphasis on capturing spatial and temporal heterogeneity (Blossfeld, 2013). The O_2_ optodes were prepared as previously described by Merl & Koren (2020). 76.5 mg of the indicator dye PtTFPP (Platinum(II)-meso(2,3,4,5,6-pentafluoro)phenyl-porphyrin, Frontier Scientific, Newark, USA), 76.8 mg of the reference dye MY (Macrolex® fluorescence yellow 10GN, Lanxess, Köln, Germany), and 5 g Polystyrene (Sigma Aldrich, Burlington, USA) were dissolved in 50 g Toluene. This ‘sensor cocktail’ was knife-coated onto a PET foil (Optimont® 501, Bleher Folientechnik GmbH, Ditzingen-Heimerdingen, Germany) using a film applicator (BYK, Wesel, Germany) and a layer of ∼10 μm was obtained after the toluene evaporated.

### Soil

The soil was collected from a long-term experimental site in Rotthalmünster, Germany (N48°21′, E13°11′). It was characterized as Haplic Luvisol with a silty loamy texture (19 % clay, 71 % silt, 10 % sand), a pH (CaCl_2_) of 6.74, with 0.13 % CaCO_3_. The soil organic carbon content was 1.21 %, total soil nitrogen was 0.14 %, and C:N ratio was 8.65. Soil was air-dried, sieved to 10 mm mesh size, and stored at 4 °C until the beginning of the experiment.

### Experimental setup

The experiment was conducted in rhizoboxes with inner dimensions of 20 cm x 2.2 cm x 40 cm (width x depth x height). Each box had three holes (7 mm diameter) in the bottom part allowing for drainage. The inner sides were lined with a polyester wick (Breatex™ 150, Fibertex Nonwovens A/S, Aalborg, Denmark) facilitating rewetting and soil moisture distribution. The “window side” of the rhizoboxes consisted of an opaque frame and transparent, removable acrylic glass windows (Linatex A/S, Herlev, Denmark) (Figure 1). The O_2_ optode was adhered onto the grid at the window side of the rhizoboxes with insulation tape. Rhizoboxes were closed using a rubber seal and C-clamps.

**Figure 1.**
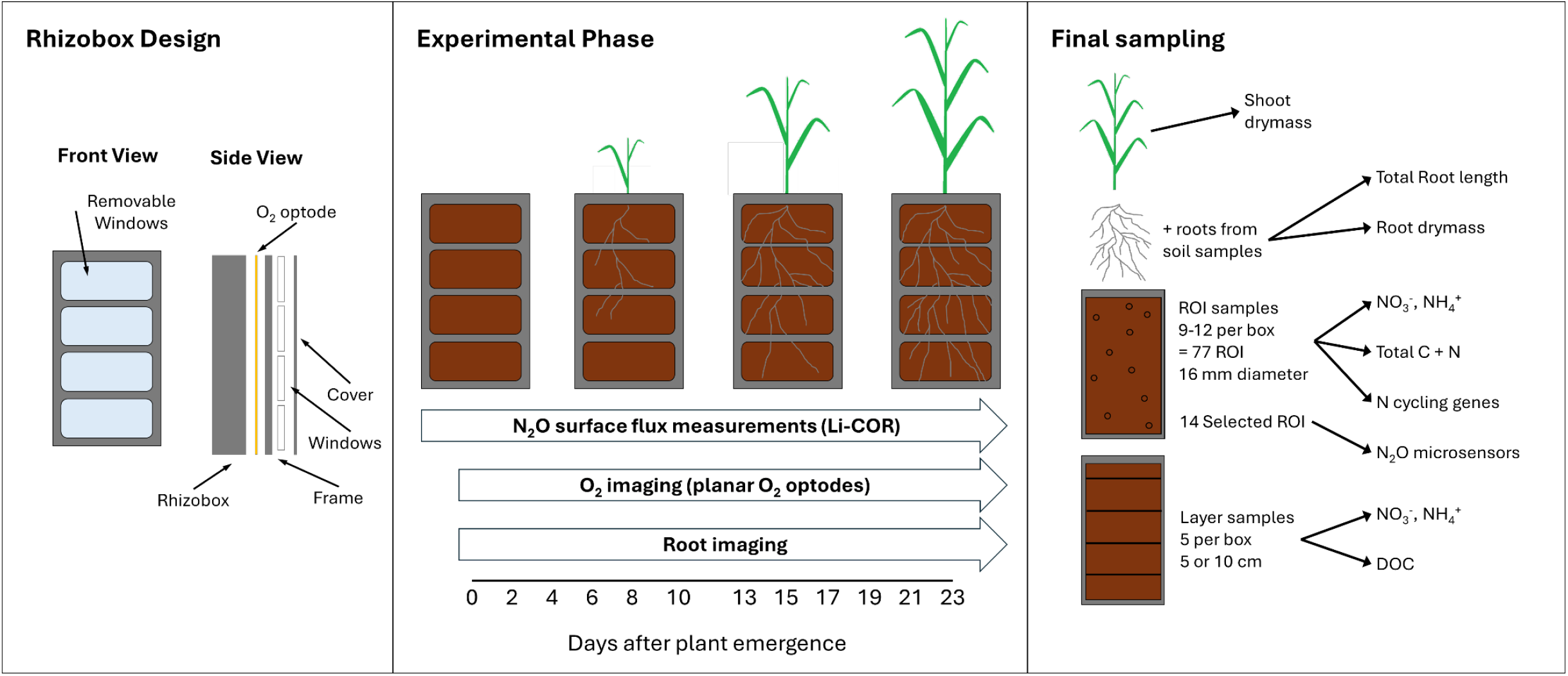
Schematic overview of rhizobox design, experimental phase, and final sampling of the experiment. The rhizobox covers consisted of an opaque frame with 4 removable window panels. To monitor O_2_ concentrations in soil, an O_2_-sensitive optode was attached to the frame before closing the rhizoboxes.

Dry soil was mixed with finely ground fertilizer (Pioner NPK Makro BLÅ, Azelis A/S, Kgs. Lyngby, Denmark) supplying 200 mg N kg^-1^, 47 mg P kg^-1^, 377 mg K kg^-1^, 64 mg S kg^-1^, and 49 mg Mg kg^-1^. Seven rhizoboxes were filled with 2.2 kg of soil per box to a height of 38 cm, resulting in a bulk density of 1.3 g cm^-3^. Soil was rewetted from the bottom over a period of 112 h and pre-incubated in a climate chamber (Weiss-Umwelttechnik GmbH, Gießen, Germany) for 10 days. One rhizobox was equipped with two FDR sensors (ECH2O 5TE, Decagon Devices, Pullman, USA) at 5-10 cm and 30-35 cm depth measuring volumetric soil water content and soil temperature. Rhizoboxes were weighed and watered every two-to-three days, always after N_2_O flux measurements and O_2_ imaging. Conditions in the climate chamber during preincubation and the experiment were as follows: 16 hours daylight with 300 μmol m^-2^ s^-1^ light intensity, 20 °C day temperature, 15 °C night temperature, 50 % relative humidity day, 60 % relative humidity night.

Maize seeds (*Zey mays* L. cv KWS Stabil) were pregerminated on wet paper for 3 days, and one germinated seed was planted into each box 10 days after rewetting the soil. However, plants emerged in only two rhizoboxes (3 and 4). From the remaining five rhizoboxes, old seeds were removed and small maize seedlings of the same variety were planted. Measurements were conducted on the same days in relation to plant age (i.e., days after emergence) in all rhizoboxes to ensure comparability across replicates.

### N_2_O surface flux analysis and calculations

Surface fluxes of N_2_O emitted from soil were measured every two-to-three days using a N_2_O/H_2_O Trace Gas Analyzer (LI-7820, Li-COR, Lincoln, USA). Rhizoboxes were put into chambers consisting of an opaque bucket and a transparent chamber top. Chamber height was 50 cm, with a total volume of 36.2 L or adjusted to 150 cm (106.9 L) according to plant size. Chamber closing times were 4 and 10 minutes for small and large chambers, respectively. The temperature in the chamber was measured once at closing time and again before opening. The mean temperature was used for flux calculations. N_2_O fluxes were calculated using the R package *goFlux* v0.2.0 (Rheault *et al*., 2024) and cumulative N_2_O emissions were calculated using *agg*.*fluxes* from the R package *gasfluxes* v0.6-2 (Fuß, 2020).

### O_2_ and root imaging setup

The imaging of the O_2_ optodes was conducted with a single-lens reflex (SLR) camera (EOS 1300D, Canon, Japan) equipped with an objective lens (EFS 18-55 mm Canon, Japan), and an orange 530 nm longpass filter (OG530 SCHOTT, 52 mm × 2 mm) with a plastic filter (#10 medium yellow, LEEfilters.com) attached in front of the long-pass filter to prevent autofluorescence of the longpass filter. For excitation of the optode, a blue LED (470 nm, r-s components, Copenhagen, Denmark) was used. The LED was controlled by a USB-controlled LED driver unit (imaging.fish-n-chips.de). The software look’RGB (imaging.fish-n-chips.de) was used to gather the images and to control the camera and LED. Calibration of the O_2_ optode was done as described in detail by (Merl & Koren, 2020). O_2_ imaging was conducted in two-to-three-day intervals, always on the same days as surface N_2_O flux measurements. Image processing and analysis was done using the software ImageJ (imagej.nih.gov/ij/) using a ratiometric approach (Larsen *et al*., 2011; Merl & Koren, 2020). For the false-color images displaying O_2_ concentration in % air saturation, the color palette *batlowK* (Scientific Color Maps v8.0.1, (Crameri, 2018) was used. We calculated the anoxic area of the rhizobox window as the proportion of the image area with an O_2_ concentration < 10 % air saturation (28.4 μmol O_2_/L) which is in the range of conditions promoting denitrification (Nakajima *et al*., 1984; Bonin *et al*., 1989) similar to previous studies (Zhu *et al*., 2015; Keiluweit *et al*., 2018).

Prior to O_2_ imaging, a photo of the rhizoboxes was taken with ‘normal light conditions’ to monitor root growth on the rhizobox window side. These root images were analyzed with RootPainter (Smith *et al*., 2022) to segment roots from the soil background. A model trained with randomly selected images was used to segment roots on all the images. Next, root length and root area were estimated from skeletonized images with RhizoVisionExplorer (Seethepalli *et al*., 2021). For image pre-processing, we used a thresholding level of 200, removed non-root objects that were smaller than 1 mm, an edge smoothing threshold of 2, and a pruning threshold of 5.

### N_2_O microsensor measurements

We used needle-shaped clark-type N_2_O microsensors that are based on an indium cathode in organic electrolyte, and an O_2_-removing compartment measuring N_2_O partial pressures (Pa) in both gas and liquid (Andersen *et al*., 2001; Revsbech, 2021). Sensors were made inhouse, similar to those commercially available from Unisense (Unisense A/S, Aarhus, Denmark). The software SensorTrace Pro (Unisense A/S, Aarhus, Denmark) was used to introduce the sensor into the soil by a computer-controlled micromanipulator and to record the sensor signals. Calibration of the N_2_O microsensors was performed in N_2_O-free water by addition of an N_2_O-Standard (27 mM N_2_O). All microsensors displayed a linear response curve during calibrations. N_2_O microsensor measurements were conducted 23 days after plant emergence. Small holes (2 × 2 mm) were cut into the optode foil and N_2_O depth profiles (0-5 mm) were measured in 500 μm steps. In total, 14 regions of interest (ROI) in three different rhizoboxes (1, 4, 5) were chosen for N_2_O microsensor measurements based on root development and O_2_ concentration.

### Harvest and soil sampling

The two rhizoboxes (Nr. 3 and 4), with initial seeds growing normally, were harvested 39 and 37 days after rewetting, corresponding to 25 and 21 days of plant growth, respectively. Rhizoboxes 1, 2, 5, 6, 7 with replanted seedlings were harvested 53, 49, 52, 50, and 52 days after rewetting, corresponding to 25, 21, 24, 22, and 24 days of plant growth. Due to the large workload of sampling and processing samples, it was not possible to harvest and sample all rhizoboxes on the same day. Shoots were cut above the surface and dried at 60 °C to determine shoot dry mass. Crown roots were added to the root samples.

In each rhizobox, regions of interest (ROI) were defined based on root development and O_2_ concentration. Soil from ROI was sampled from the window side of the rhizoboxes with small aluminum cylinders (16 mm inner diameter, 10 mm depth) (UGT Umwelt-Geräte-Technik GmbH, Müncheberg, Germany) and the roots were carefully separated from the soil. The soil was then homogenized and divided into three subsamples: ∼1 g soil (dry-weight-equivalent) was frozen in Eppendorf tubes (at – 20 °C for analysis of bacterial functional genes, ∼1 g soil (dry-weight-equivalent) was stored in 10 ml centrifuge tubes at 4°C for analysis of mineral N, and the remaining soil (ca. 1-1.2 g) was dried at 105 °C to determine soil water content and total C and N content.

Additional soil samples were taken from rhizoboxes in 5 depths: 0-5 cm, 5-10 cm, 10-20 cm, 20-30 cm, 30-38 cm using a cork borer as auger (inner diameter 2 cm). For each depth layer, eight samples were taken and homogenized into one composite sample, after careful removal of the roots. Each composite sample was then divided into three subsamples: ∼50 g soil (dry-weight-equivalent) was stored at 4°C for analysis of mineral N, ∼10 g soil (dry-weight-equivalent) was stored at 4°C for analysis of water-extractable C_org_ (WEOC), and ∼ 5 g soil (dry-weight-equivalent) were dried at 105° C to determine soil water content. Roots were washed from the remaining soil and stored in 30 % ethanol at 4°C.

### Root scanning and analysis

Roots were scanned on a flatbed scanner (Epson, Suwa, Japan) and the software RhizoVisionExplorer (Seethepalli *et al*., 2021) was used to extract total root length and total root volume from the scanned images. For image pre-processing, we used a thresholding level of 200, removed non-root objects that were smaller than 0.1 px, an edge smoothing threshold of 2, and a pruning threshold of 5. Root diameter classes were classified as <1mm, 1-2 mm, and >2 mm. Total root dry weight was determined by drying roots at 105 °C. We refer to root length at the rhizobox window estimated with root imaging as *planar root length* and root length at the final harvest measured with root scanning as *total root length* (Wacker *et al*., 2024).

### Soil analysis

Soil mineral N was extracted by shaking the subsample with 1 *M* KCl (ratio 1:5 w:v) on an overhead shaker with 40 rpm for 60 min, filtered (Whatman Grade 42 for depth layer soil samples, 0.8 μm CA syringe filters for ROI samples) and analyzed colorimetrically on a continuous flow analyzer (AA3, Seal Analytical, Norderstedt, Germany). WEOC was extracted by thoroughly shaking soil with ultrapure water (ratio 1:2 w:v) for 60 s, filtered through 0.45 μm PES syringe filters, and analyzed on an elemental analyzer (multi N/C 2100 S, Analytik Jena, Jena, Germany). Total C and N content in ROI samples was analyzed on an elemental analyzer (Vario EL III, Elementar, Langenseibold, Germany).

### DNA extraction and quantitative PCR

DNA was extracted from 400 mg of freeze-dried soil for each sample using the NucleoSpin Soil kit (Macherey-Nagel, Düren, Germany), following the manufacturer’s instructions. DNA quality was checked using a NanoDrop (Thermo Fisher Scientific, Waltham, MA, USA), before quantification on a Qubit fluorimeter using the Broad Range double stranded DNA kit (Thermo Fisher Scientific). The abundance of total archaea and bacteria (16S rRNA gene), denitrifiers (*nirK* and *nirS*) and N_2_O reducers (*nosZ*I and *nosZ*II) were determined using real-time quantitative PCR (qPCR) using specific primers for each gene. The qPCR reactions were performed in two independent runs in a reaction volume of 15 μL containing iQ™ SYBR Green Supermix (Bio-Rad, Hercules, CA, USA), 0.1 % bovine serum albumin (New England Biolabs, Ipswich, MA, USA), primers and 2 ng of template DNA on a CFX Connect Real-Time System (Bio-Rad). Primers, qPCR conditions, and amplification efficiencies are presented in Supplementary Table S1. Standard curves were generated by serial dilutions of linearized plasmids containing a fragment of the specific gene. The amplifications were validated by melting curve analyses and agarose gel electrophoreses. Potential inhibition of PCR reactions was initially checked by amplifying a known amount of the pGEM-T plasmid (Promega, Madison, WI, USA) with the plasmid specific M13F/M13R primer set (Supplementary Table S1) and 2 ng of DNA template or non-template controls for each sample. No inhibition was detected with the amount of DNA used.

### Calculations and statistics

All calculations and statistics were carried out using the statistical software R v4.3.3 (R Core Team, 2024). Simple linear regressions were applied to identify relationships between soil moisture, planar root length, or anoxic soil area. Similarly, we applied simple linear regression models to analyze relationships between soil N_2_O concentrations, O_2_ concentrations, N availability, and abundance of N cycling genes in ROI at the end of the experiment. To identify the factors driving N_2_O fluxes, we applied linear mixed effect models (lme) using the lme function from the package nlme v3.1-131 (Pinheiro *et al*., 2017). We tested models for the experimental period with and without plant growth, respectively, as well as for the whole experimental period. To account for repeated measurements, replicates were set as random effect. Models were compared using maximum likelihood (ML), selected using AIC (Akaike’s information criterion), and fitted using restricted maximum likelihood (REML). Pseudo-*R*^2^ for lme was calculated using r.squaredGLMM from the package MuMIn v1.42.1 (Barton, 2018).

## Results

### Plant and root growth

Root development followed the same pattern in all rhizoboxes. Planar root length increased slowly during the first week after emergence, then increased almost linearly until harvest (Fig. 2 a). Total root length at harvest was 161.4 ± 48.3, root dry weight was 1.11 ± 0.47 g, and root dry weight was positively correlated with total root length (*R*^2^=0.83, *p* < 0.01). Maize shoot dry weight at harvest 4.24 ± 1.16 g, and root:shoot ratio was 0.25 ± 0.03 (Supplementary Table S2). Shoot N content was 3.5 ± 0.4 %.

**Figure 2.**
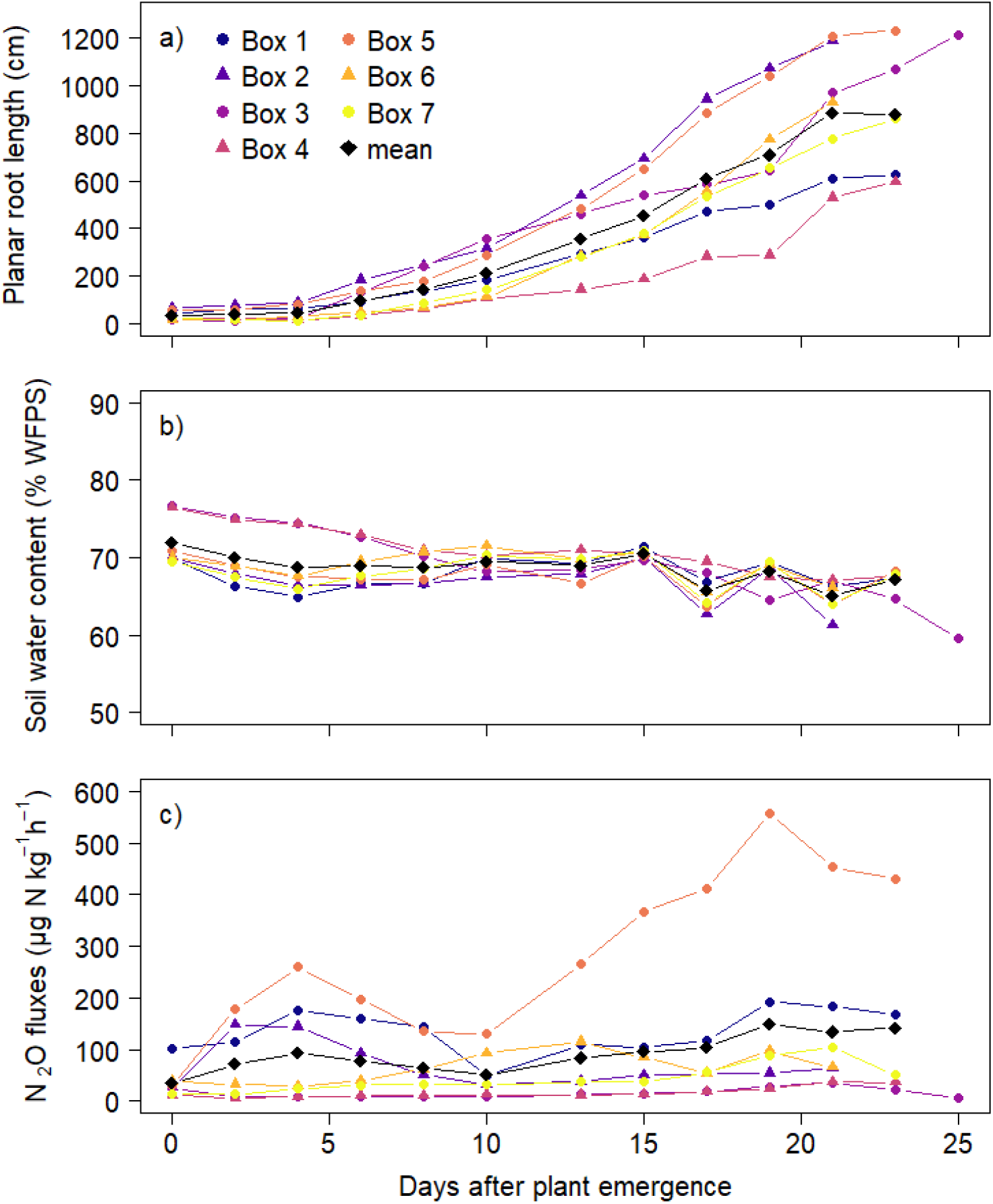
(a) Planar root length (cm), (b) soil water content (% water-filled pore space), and (c) N_2_O fluxes (μg N_2_O-N kg^-1^ h^-1^). Colored points and triangles represent each individual rhizobox, black diamonds represent means (n=7).

### Soil Moisture

After rewetting, soil moisture ranged between 84 and 89 % WFPS (Supplementary Fig S1) and was kept at 65-70 % WFPS during plant growth, providing favorable conditions for anaerobic processes such as denitrification (Fig. 2b). Soil moisture sensors showed that soil moisture was very stable in the 30-35 cm layer and fluctuated most in the top 5-10 cm layer (Supplementary Figure S2).

### N_2_O fluxes and cumulative emissions

The measured N_2_O fluxes showed large variations between rhizoboxes (Fig. 2c, Supplementary Fig. S1). During the acclimation phase, N_2_O fluxes were comparable in all rhizoboxes and decreased for the first two weeks after rewetting. In the rhizoboxes that had to be replanted, N_2_O fluxes remained relatively stable until plant emergence (Supplementary Fig S1). In rhizoboxes 1, 2, and 5, N_2_O fluxes increased directly after plant emergence, in rhizobox 6, N_2_O fluxes increased slightly a week after plant emergence, and N_2_O fluxes from rhizoboxes 1, 5, and 7 increased towards the end of the experiment (Fig. 2c). N_2_O fluxes from box 1 were higher than all other rhizoboxes before plant emergence (Supplementary Fig. S1), and highest from rhizobox 5 after plant emergence (Fig. 2c).

Cumulative N_2_O emissions during the planted phase ranged from 8.5 N_2_O-N kg^-1^ to 157.5 and (Supplementary Fig. S3).

### Soil O_2_ concentrations and anoxic soil fraction

O_2_ imaging started 10 days after rewetting in the rhizoboxes, and most of the optode-soil interface was anoxic at this time (Example for Rhizobox 1 in Figure 3, other data found in Supplementary Figure S4 and S5). Overall, O_2_ concentrations increased over time in all depths, although watering also led to formation of anoxic conditions in the upper soil layers without root growth (Figure 3). In the deeper soil layers, the rhizobox window area covered with the O_2_-sensitive planar optode showed almost completely anoxic conditions throughout the experiment. During plant growth, the anoxic fraction decreased in all layers in all rhizoboxes as the soil became increasingly more oxic. O_2_ concentrations rapidly increased in the topmost soil layer where more than 90 % of the window area was oxic 24 days after plant emergence. O_2_ concentrations in the depths below followed similar patterns, with a time delay. With increasing root growth, larger areas became oxic, but in the direct vicinity of roots, O_2_ concentrations were lower compared to surrounding bulk soil.

**Figure 3.**
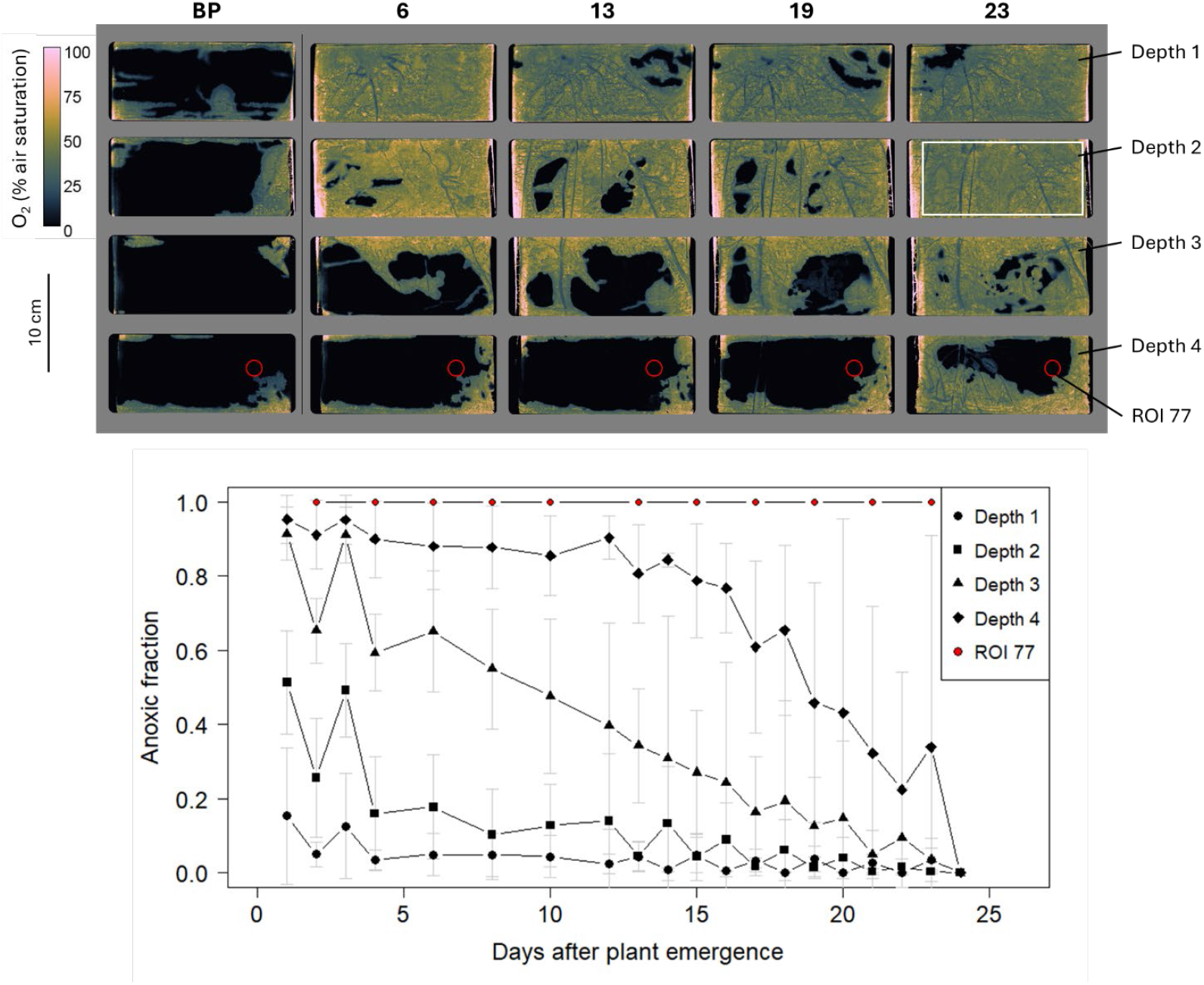
Top: False color images of the O_2_ concentrations in Rhizobox 1 before plant emergence (BP) and on Day 6, 13, 19, and 23 after plant emergence measured with planar O_2_ optodes. The four ‘windows’ of the rhizobox correspond to the four depths. The white rectangle is exemplary for the area used to calculate the anoxic fraction of each depth. Lighter areas outside the white rectangle are artifacts at the edges of the optode and were not included in calculations. The red circle represents ROI 77. Bottom: Fraction of anoxic area (< 10% O_2_ air saturation = 2.1 % O_2_) in all rhizoboxes. Different symbols represent the four depths in the rhizoboxes and ROI 77. Mean ± standard deviation for n=7.

### Interactions between root growth, soil moisture, soil profile O_**2**_ **concentrations, and surface N**_2_O fluxes

Soil water content and mean anoxic fraction were both negatively correlated with root length (*R*^2^=0.27 and 0.85, respectively, *p* < 0.0001), while the anoxic fraction was positively correlated with soil water content (*R*^2^=0.42, *p* < 0.0001) (Table 1, Supplementary Figure S6). Linear regressions between surface N_2_O fluxes and either soil water content, root length, or anoxic fraction yielded only weak relationships (Table 1, Supplementary Figure S6).

**Table 1.** Results of regression analyses (coefficients of determination (Adjusted *R*^2^), *p*-values, and sample size *n*) of the relationship between soil water content, root length, anoxic surface fraction, and N_2_O surface fluxes (ln transformed) in rhizoboxes.

| Model | $n$ | $R^2$ | $p$-value |
| --- | --- | --- | --- |
| *Simple linear regressions* |  |  |  |
| Soil water content ~ Root length | 76 | 0.2738 | $7.42 \times 10^{-07}$ |
| Anoxic fraction ~ Soil water content | 76 | 0.4247 | $1.099 \times 10^{-10}$ |
| Anoxic fraction ~ Root length | 76 | 0.8482 | $< 2.2 \times 10^{-16}$ |
| Root dry weight ~ total root length | 76 | 0.8303 | 0.002693 |
| Soil water content ~ total root length | 76 | 0.8899 | 0.0008959 |
| $\ln(N_2O) \sim$ Root length | 76 | 0.06972 | 0.01263 |
| $\ln(N_2O) \sim$ Soil water content | 76 | 0.1058 | 0.002411 |
| $\ln(N_2O) \sim$ Anoxic fraction | 76 | 0.115 | 0.001593 |
| *Linear mixed effect model* |  |  |  |
| $\ln(N_2O) \sim$ Soil water content * Anoxic Fraction<br>Random effect: Rhizobox Nr (replicate) | 76 | 0.857 (cond.)*<br>0.091 (marg.)* | 0.0097 |

\*For the linear mixed effect model (lme), the conditional $R^2$ is the variance explained by the entire model, including both fixed and random effects, while the marginal $R^2$ represents only the variance explained by the fixed effects.

The best model explaining surface N_2_O fluxes included an interaction between soil water content and anoxic fraction as fixed factors and the rhizobox as random factor (*p* < 0.01). At a high anoxic fraction, ln(N_2_O) was negatively correlated with soil water content, while ln(N_2_O) was positively correlated with soil water content for low anoxic fractions pointing towards interactive effects of soil water content and anoxia on N_2_O formation and reduction. The conditional *R*^2^ of the full model was 0.86, while the marginal *R*^2^ of the model with only the fixed effect (i.e., without the random factor) was 0.09 highlighting the large differences between rhizoboxes (Table 1, Supplementary Figure S7). Results of regression analyses for the dataset covering the whole experimental period can be found in the Supplementary Table S3.

### Soil analyses (total CN, mineral N, WEOC)

Soil analyses showed similar depth patterns for analysis of soil layers and ROI (Figure 4). Soil NO_3_^−^ content decreased with depth, from 320-1299 mg N kg^-1^ in the upper 5 cm to 1.7-502 mg N kg^-1^ in the 5-10 cm layer to 0.04-123 mg N kg^-1^ below 10 cm (Figure 4 a). Soil NH_4_^+^ contents were much lower compared to NO_3_^−^ content. NH_4_^+^ contents were lowest in the 5-20 cm layers and ranged between 0 and 45 mg N kg^-1^ in the others (Figure 4b). Total soil N in ROI ranged between 0.09 and 0.26 % without clear differences in different depths (Supplementary Figure S8). Total soil C in ROI mostly ranged between 1.11 and 1.25 % which was similar to the initial soil C_org_ content (1.21 %). From 77 measured ROI, eight ROI had higher total C contents between 1.5 and 2.3 % which likely included small pieces of roots or CaCO_3_. DOC content ranged between 24.5 and 188. 7 mg C kg^-1^ with highest values in 0-5 cm layers (Figure 4c).

**Figure 4.**
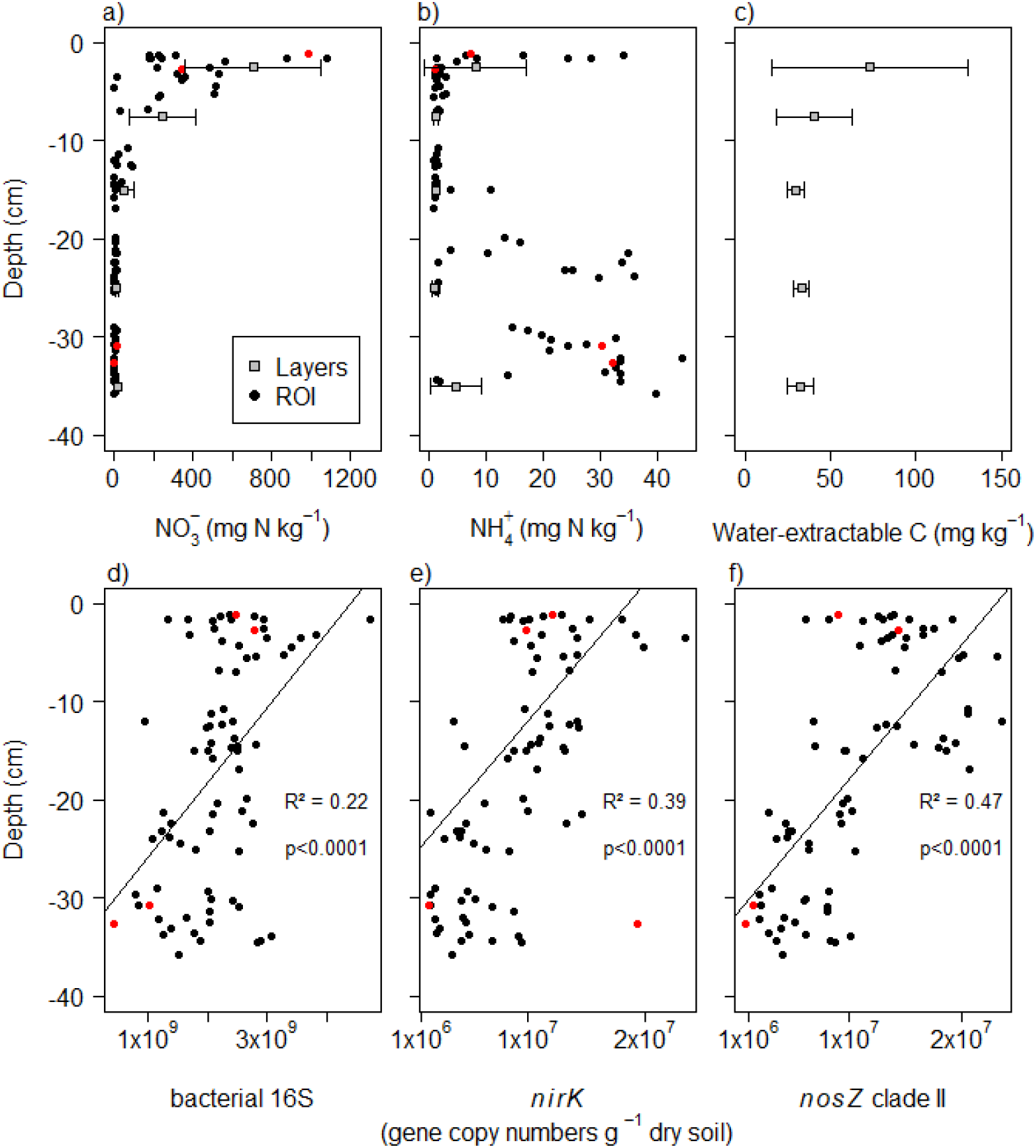
Depth distribution of (a) NO_3_^-^, (b) NH_4_^+^, and (c) DOC content in soil layers and ROI, and (d-f) depth distribution of gene abundances in ROI. Each point represents one of 77 ROI, while grey squares represent mean values ± standard deviation for different soil layers (n=7). Red circles represent ROI Nr. 1, 6, 45, and 77. For gene abundances, regression lines, adjusted R^2^, and *p*-values of simple linear regressions with depth are depicted.

### Functional genes involved in N cycling

Abundance of 16S rRNA genes, a proxy for total archaea and bacteria, decreased with increasing sampling depth (*R*^2^=0.22, *p* < 0.0001, Figure 4d, Table 2). Similarly, abundance of *nirK* and *nosZ* clade II genes decreased with increasing sampling depth (*R*^2^=0.39 and 0.47, respectively, *p* < 0.0001). For *nirS* and *nosZ* clade I, only very weak correlations with sampling depth were found (Table 2). Except for *nirS*, gene abundances were negatively correlated with soil NH_4_ ^+^ concentrations (Supplementary Table S4). However, NH_4_ ^+^ concentrations were also negatively correlated with soil depth. The ratio of denitrifiers (*nirK* + *nirS*) to N_2_O reducers (*nosZ*I + *nosZ*II) increased with increasing NO_3_ ^-^ content in ROI (*R*^2^=0.2, *p* < 0.0001).

**Table 2.** Results of simple linear regression analyses (coefficients of determination (Adjusted *R*^2^), *p*-values, and sample size *n*) of the relationship between N_2_O, O_2_, N concentrations, root age, soil depth, and abundance of nitrogen cycling genes in ROI. Significant relationships (*p* < 0.05) are highlighted in bold.

| Model | $n$ | $R^2$ | $p$-value |
| --- | --- | --- | --- |
| **Min. $\text{N}_2\text{O}$ ~ $\text{O}_2$ max** | **14** | **0.5304** | **0.001893** |
| **Max. $\text{N}_2\text{O}$ ~ N concentration** | **14** | **0.5227** | **0.002097** |
| **16S ~ Depth** | **77** | **0.2193** | **$1.04 \times 10^{-05}$** |
| ***nirK* ~ Depth** | **77** | **0.3882** | **$8.65 \times 10^{-10}$** |
| *nirS* ~ Depth | 77 | 0.04388 | 0.03745 |
| *nosZI* ~ Depth | 77 | 0.07119 | 0.01085 |
| ***nosZII* ~ Depth** | **77** | **0.4691** | **$3.891 \times 10^{-12}$** |
| **$\text{NH}_4^+$ ~ Depth** | **77** | **0.3325** | **$2.436 \times 10^{-08}$** |
| ***nirK* ~ root age** | **77** | **0.2431** | **$3.127 \times 10^{-06}$** |
| ***nosZII* ~ root age** | **77** | **0.1822** | **$6.433 \times 10^{-05}$** |
| **$(\text{nirK}+\text{nirS}) / (\text{nosZI}+\text{nosZII}) \sim \text{NO}_3^-$** | **76** | **0.2** | **$3.045 \times 10^{-05}$** |

### N_2_O microsensor profiles in ROI and their relationship with influencing parameters

N_2_O concentrations measured with N_2_O microsensors in ROI at the end of the experiment showed very high variability in N_2_O concentrations (Figure 5, Supplementary Figure S9). N_2_O concentrations ranged between 0 and 100 μmol N_2_O L^-1^ with highest N_2_O concentrations measured in ROI 1 close to the shoot and its crown roots in 1.1 cm depth and second highest measured in ROI 77 around a small root at the bottom of the rhizobox at 30.8 cm depth.

**Figure 5.**
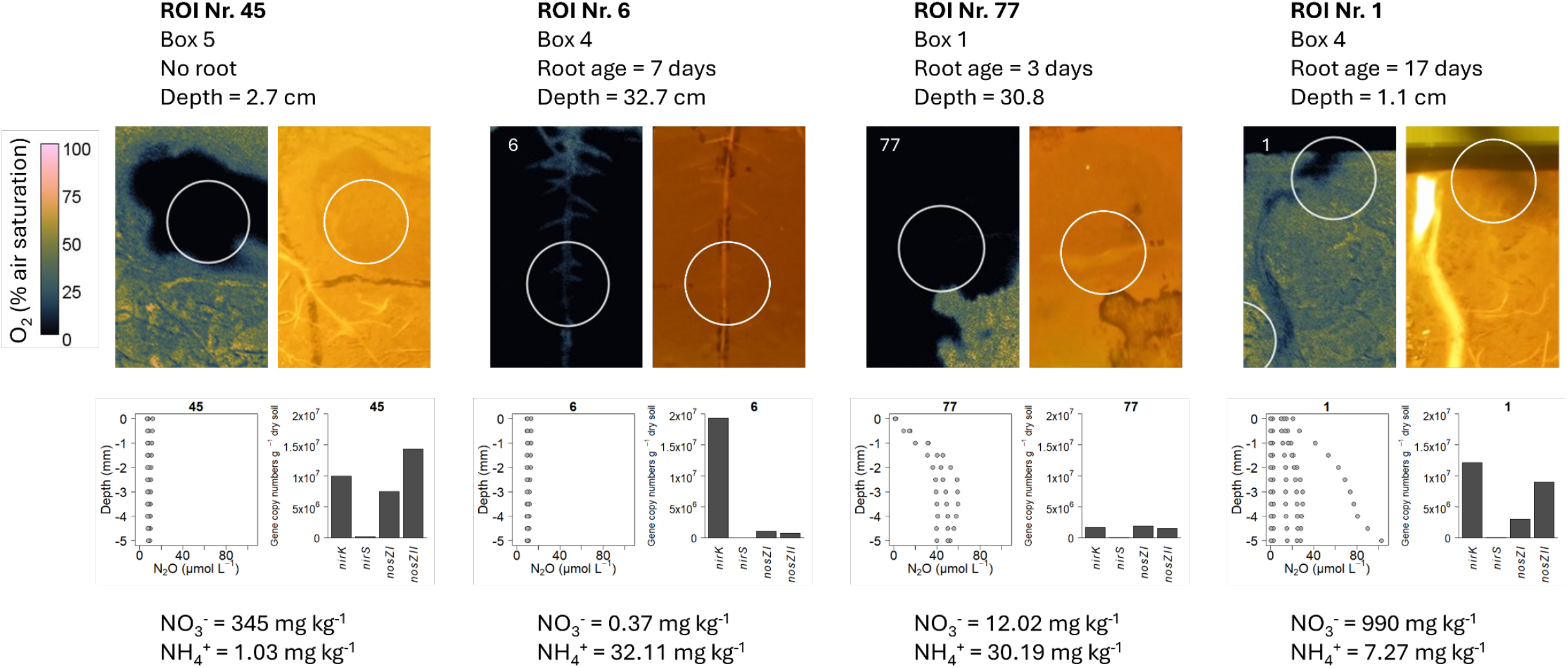
Characterization of four exemplary ROI at the end of the experiment. False-color image showing O_2_ concentration, photo showing root development, N_2_O microsensor depth profiles (μmol N_2_O), barplot showing gene abundances (gene copy numbers g^-1^ dry soil), and mineral N content (mg N kg^-1^). ROI diameter = 16 mm. Root age refers to the days of root growth in the respective ROI.

For most ROI, changes in N_2_O concentration within the same measurement profile were in the range of a few μmol (Supplementary Figure S9). There were exceptions, as ROI 1 where one N_2_O profile increased from 20 to 100 μmol L^-1^ and ROI 77 where N_2_O concentrations increased from 0 to 40-60 μmol L^-1^ in 2-5 mm depth (Figure 5). Minimum N_2_O concentrations in 500 μm depth (i.e, in direct proximity to the optode) were negatively correlated with maximum O_2_ concentration (*R*^2^=0.53, *p* < 0.01, Table 2, Supplementary Figure S10), while maximum N_2_O concentrations in 500 μm depth were positively correlated with total N availability in all ROI (*R*^2^=0.52, *p* < 0.01, Table 2, Supplementary Figure S10).

## Discussion

### Microsite N_2_O formation depended on root growth, O_2_ and N availability

N_2_O microsensor measurements showed very high spatial heterogeneity of N_2_O formation in soil. Minimum N_2_O concentrations were higher in ROI with lower O_2_ concentrations confirming that O_2_ availability was a major control of denitrification at the microsite scale. Oxygen concentration is one of the most important controls of denitrification as it determines when microorganisms switch to anaerobic respiration (Groffman *et al*., 1988; Schlüter *et al*., 2024). This, however, does not ascertain that denitrification will occur, as nitrate can be limiting (Smith & Tiedje, 1979; Hojberg *et al*., 1994; Schlüter *et al*., 2024), which has been documented in relation to plant growth with rapid N uptake (von Rheinbaben & Trolldenier, 1984; Haider *et al*., 1985; Rummel *et al*., 2021). Accordingly, we found higher N_2_O concentrations in ROI with higher total N availability indicating that, under O_2_ limiting conditions, N availability is crucial to determine the magnitude of N_2_O emissions. ROI with the highest N_2_O concentrations were in the direct vicinity of roots (i.e., ROI Nr. 1, 77). In contrast, ROI 45 was characterized by high NO_3_ ^-^ content and anoxic conditions, but no roots, and only low N_2_O concentrations confirming the importance of roots on formation of N_2_O hotspots through C exudation, root and rhizosphere microbial respiration, verifying our second hypothesis. Although functional gene abundances and N_2_O concentrations were not significantly correlated in ROI, we found a positive relationship between the (*nirK*+*nirS*)/(*nosZI*+*nosZII*) ratio and NO_3_ ^-^ availability indicating higher relative genetic potential for N_2_O reduction at lower soil NO_3_ ^-^ content. This relationship was driven by the abundance of *nosZ* clade II nitrous oxide reducers, in line with previous reports (Xu *et al*., 2020; Jones *et al*., 2022).

### Roots determined biotic and abiotic gradients in soil

Plant and root growth affected soil moisture, anoxic soil fraction, and distribution of nutrients controlling the abundance of microorganisms. Root growth determined soil moisture through root water uptake as demonstrated by significant negative correlations between soil water content and planar root length throughout the experiment and between soil water content and total root length at final harvest. Further, soil moisture sensors showed that soil moisture was constant at the bottom of the rhizoboxes, while fluctuating more in the upper soil layer with higher root density. Using planar optodes, we were able to show increasing O_2_ concentrations with increasing plant and root growth. Especially, roots growing into deeper soil layers led to increasing O_2_ concentrations at the optode-soil interface. We anticipate a combination of several explanations: (1) plant roots can increase the diffusion of O_2_ through the soil (Jensen & Kirkham, 1963), (2) maize roots form aerenchyma under O_2_-limited conditions (≤ 3% O_2_ ≈ 14.3 % O_2_ air saturation (Gunawardena *et al*., 2001) facilitating transport of O_2_ into anoxic soil layers, (3) and roots may have ‘pushed’ the optode away from the soil surface in some spots creating air-filled cavities between the optode and the soil (Merl *et al*., 2023). Although roots generally increased aeration and O_2_ concentrations in soil, roots were always characterized by lower O_2_ concentrations compared to the surrounding soil confirming our first hypothesis. Most of the added NO_3_ ^-^ fertilizer was recovered in the upper 10 cm of the rhizoboxes likely due to a combination of initial rewetting the soil in the rhizoboxes from the bottom and evaporation on the soil surface causing a constant upward movement of soil water, transporting NO_3_ ^-^ to the upper soil layer throughout the experiment. Root growth increased availability of organic C through rhizodeposition leading to higher DOC concentrations in upper soil layers with higher root density. Although root exudation per root surface area decreases with growth/age, total exuded C rates increase with increasing root growth (Santangeli *et al*., 2024). Especially, maize crown/brace roots can exude large quantities of mucilage containing high amounts of sugars (Werner *et al*., 2022) contributing to high availability of DOC in the uppermost soil layers. Higher availability of DOC and NO_3_ ^-^ in upper soil layers promoted microbial growth as indicated by decreasing abundances of 16S rRNA, *nirK* and *nosZII* genes with soil depth. Abundances of *nirK* and *nosZII* genes also increased with increasing root age confirming the influence of root growth on microbial denitrifiers. Abundances of *nirS* and *nosZI* were much lower compared to abundances of *nirK* and *nosZII*, respectively, and were not affected by nutrient concentrations or root growth indicating niche differentiation between *nirK-* and *nirS-*type denitrifiers and *nosZ* clade I and II nitrous oxide reducers, respectively (Jones & Hallin, 2010; Achouak *et al*., 2019; Wang *et al*., 2024).

### Soil moisture and anoxic fraction controlled N_2_O fluxes

N_2_O fluxes showed distinct patterns with initially high fluxes decreasing until about two weeks after rewetting the soil, followed by a period with lower fluxes. Emergence of plants stimulated N_2_O production leading to clear increases in N_2_O fluxes within a week following plant emergence. However, correlations between N_2_O fluxes and planar root length or cumulative N_2_O emissions and total root length at harvest did not yield significant relationships as we would have expected. Nonetheless, our study confirmed intercorrelation of root growth and N_2_O formation as N_2_O fluxes were controlled by an interaction of the anoxic fraction and soil moisture, confirming interacting effects of root water uptake and anaerobicity on N_2_O formation and reduction. For low anoxic fractions, increasing soil moisture led to increasing N_2_O fluxes, while for high anoxic fractions higher soil moisture led to lower N_2_O fluxes due to the reduction of N_2_O to N_2_ (Davidson, 1991; Rohe *et al*., 2021). In addition, differences between replicates explained most of the variance in N_2_O fluxes. Soil heterogeneity is known to strongly control the formation of denitrifying N_2_O hotspots causing very high variation within replicates of the same treatment (Folorunso & Rolston, 1984; Christensen *et al*., 1990; Schlüter *et al*., 2024). Plant and root growth further increased heterogeneity in soil but also differences between replicates.

### The uppermost soil layers contributed largely to N_2_O fluxes

We anticipated that some zones of the rhizoboxes were more likely to contribute to N_2_O formation – the uppermost and the lowest soil layers. The upper 5 cm soil layer was characterized by very high NO_3_ ^-^ concentrations (up to >1000 mg NO_3_ ^-^-N kg^-1^), highest concentrations of DOC, and highest abundances of denitrifying and N_2_O reducing genes (*nirK* and *nosZII*). Although O_2_ concentrations in the uppermost soil layer increased fast, small anoxic areas were identified with the O_2_ optodes at the later/last measurement days. Furthermore, rewatering from the top kept soil continuously wet which (together with high NO_3_ ^-^ and DOC) provided optimal conditions for denitrifiers as confirmed by microsensor measurements (ROI 1). Short distance to the soil surface facilitated N_2_O diffusion to the surface and its loss to the atmosphere. Accordingly, cumulative N_2_O emissions were positively correlated with soil NO_3_ ^-^ content in the uppermost soil layer. In contrast, the bottom layer of the rhizoboxes had larger areas that remained anoxic throughout the experiment. NO_3_ ^-^ concentrations in the bottom layer were low (15 mg NO_3_ ^-^-N kg^-1^, 200 μM NO_3_ ^-^), but sufficient to sustain denitrifying microorganisms (Palmer & Horn, 2015) as confirmed by N_2_O microsensor profiles indicating substantial N_2_O formation around a small root in ROI 77. Continuous high soil moisture and long diffusion pathways from the lower layers of the rhizobox to the soil surface likely promoted N_2_O reduction to N_2_, which has been reported for this particular soil (Malique *et al*., 2019; Rummel *et al*., 2021; Yankelzon *et al*., 2024a).

## Conclusions

We provide the first *in-situ* measurements of N_2_O concentration profiles showing distinct patterns with highest N_2_O concentrations around roots compared to bulk soil. Our study confirmed that O_2_ and N availability control N_2_O production at the process scale while emphasizing the importance of roots in shaping N_2_O hotspots. Plant roots determined depth gradients in the soil of the rhizoboxes with higher DOC and lower soil moisture in the upper soil layers. Abundances of total bacteria, denitrifying *nirK* and N_2_O reducing *nosZII* responded to resource availability with higher abundances in upper soil layers with high nutrient contents and more root growth. We argue that these uppermost soil layers largely contributed to N_2_O formation and N_2_O fluxes measured at the soil surface. Combining O_2_ optode and root imaging proved to be a successful approach to monitor major controls of rhizosphere N cycling and N_2_O formation at high spatial and temporal resolution. Carefully choosing ROIs allowed reducing the number of samples, while covering a broad range of soil conditions revealing correlations between N_2_O concentrations, substrate availability, and microbial gene abundances.

## Acknowledgements

The authors would like to thank Lars Borregaard Pedersen for excellent technical support, Tomke S. Wacker, Frederik van der Bom, and Lorène Siegwart for support on root imaging, scanning, and analysis, Andrey Kalinichev and Fabian Steininger for help with optode calibration, Sabine Rautenberg and Claudia Kuntz for soil analyses, Franziska Eller for the climate chambers and LiCOR, and Per Lennart Ambus and Kristian Thorup-Kristensen for feedback on the experimental setup.

PSR was supported by a Novo Nordisk Fonden (NNF) Postdoctoral Fellowship (grant number NNF22OC0079335). MRR was supported by the Pioneer Center for Research in Sustainable Agricultural Futures (Land-CRAFT), DNRF grant number P2.

## Competing interests

The authors declare no competing interests.

## Author contributions

Conceptualization: PSR, CWM, KK; Investigation: PSR, MRR, AS; Data analysis (soil, root, N_2_O): PSR; Data analysis (optodes): PSR, TM; Data analysis (qPCR): AS; Writing – original draft preparation: PSR; Writing – review and editing: MRR, AS, TM, SH, CWM, KK. All authors have read and agreed to the published version of the manuscript.

## Data availability

The data that support the findings of this study will be made openly available at http://doi.org/10.5281/zenodo.15537545.

## Notes

### Competing Interest Statement

The authors have declared no competing interest.

### Summary of Updates

Figures updated, description of methods revised

