## Supplementary for "Maize root growth, Oxygen and N availability drive formation of N_2_O hotspots in soil"

**Table S1:** Primers and thermal cycling conditions for quantification of 16S rRNA, denitrification and nitrate ammonification. Melting curve conditions are indicated in italic.

| **Process** | **Gene** Primer names | **Sequences** (5'-3') | **Conc. (µM)** | **Thermal cycling** | **Efficiency** |
|---|---|---|---|---|---|
| **Inhibition test** | M13F | GTAAAACGACGGCCAG | 0.25 | (95°C, 5 min) x 1 | / |
| | M13R | CAGGAAACAGCTATGAC | 0.25 | (95°C, 15 s; 55 °C, 30 s; 72°C, 30 s; 80°C, 5 s) x 35 | |
| | **16S rRNA**[1] | | | (95°C, 5 min) x 1 | |
| | 515F | GTGYCAGCMGCCGCGGTAA | 0.5 | (95°C, 15s, 50°C 30s, 72°C 30s, 78°C 5s) x 35 | 85-86% |
| | 926R | CCGYCAATTYMTTTRAGTTT | 0.5 | *(95°C, 15 s; (60 to 95° C, 5 s, increment 0.5°)), x 1* | |
| **Denitrification** | ***nirK***[2] | | | (95°C, 5 min) x 1 | |
| Nitrite reduction | 876F | ATYGGCGGVAYGGCGA | 0.5 | (95°C, 15 s; (63°C – 58°C, -1°/cycle), 30 s; 72°C, 30 s) x 6 | 96-100% |
| | 1040R | GCCTCGATCAGRTTRTGGTT | 0.5 | (95°C, 15 s; 58°C, 30 s; 72°C, 30 s; 80°C, 5 s) x 35 | |
| | | | | *(95°C, 15 s; (60 to 95° C, 5 s, increment 0.5°)), x 1* | |
| **Denitrification** | ***nirS***[3] | | | (95°C, 5 min) x 1 | |
| Nitrite reduction | cd3aFm | AACGYSAAGGARACSGG | 0.5 | (95°C, 15 s; (65°C – 60°C, -1°/cycle), 30 s; 72°C, 35 s) x 6 | 87-89% |
| | R3cdm | GASTTCGGRTGSGTCTTSAYGAA | 0.5 | (95°C, 15 s; 60°C, 30 s; 72°C, 35 s; 80°C, 5 s) x 35 | |
| | | | | *(95°C, 15 s; (60 to 95° C, 5 s, increment 0.5°)), x 1* | |
| **Denitrification** | ***nosZ*I**[4] | | | (95°C, 7 min) x 1 | |
| Nitrous oxide reduction | 1840F | CGCRACGGCAASAAGGTSMSSGT | 0.8 | (95°C, 15 s; (65°C – 60°C, -1°/cycle), 30 s; 72°C, 30 s) x6 | 93-94% |
| | 2090R | CAKRTGCAKSGCRTGGCAGAA | 0.8 | (95°C, 15 s; 60°C, 30 s; 72°C, 30 s; 80°C, 5 s) x 35 | |
| | | | | *(95°C, 15 s; (60 to 95° C, 10 s, increment 0.5°)), x 1* | |
| **Denitrification** | ***nosZ*II**[5] | | | (95°C, 7 min) x 1 | |
| Nitrous oxide | nosZII-F | CTIGGICCIYTKCAYAC | 0.8 | (95°C, 15 s; 54°C, 30 s; 72°C, 30 s; 77°C, 5 s) x 40 | 88-89% |
| reduction | nosZII-R | GCIGARCARAAITCBGTRC | 0.8 | (95°C, 15 s; (60 to 95° C, 10 s, increment 0.5°)), x 1 | |

[1]: Parada et al., 2016, Environ. Microbiol., https://doi.org/10.1111/1462-2920.13023 & Quince et al., 2011, BMC Bioinf., https://doi.org/10.1186/1471-2105-12-38

[2]: Henry et al., 2004, J. Microbiol. Methods, https://doi.org/10.1016/j.mimet.2004.07.002

[3]: Throbäck et al., 2004, FEMS Microbiol. Ecol., https://doi.org/10.1016/j.femsec.2004.04.011

[4]: Henry et al., 2006, Appl. Environ. Microbiol., https://doi.org/10.1128/AEM.00231-06

[5]: Jones et al., 2013, ISME J., https://doi.org/10.1038/ismej.2012.125

**Table S2:** Shoot dry weight, root dry weight, root:shoot ratio, total root length, shoot N content, and shoot N uptake for each individual rhizobox.

| Rhizobox Nr. | Shoot dry weight (g) | Root dry weight (g) | Root:Shoot ratio | Total root length (m) | N content (%) | Total shoot N content (mg) |
|---|---|---|---|---|---|---|
| 1 | 3.38 | 0.7410 | 0.22 | 102.50 | 3.1 | 104.4 |
| 2 | 4.09 | 0.9970 | 0.24 | 167.73 | 3.7 | 152.1 |
| 3 | 7.13 | 2.0480 | 0.29 | 238.88 | 2.8 | 198.7 |
| 4 | 3.03 | 0.7080 | 0.23 | 114.38 | 3.8 | 114.9 |
| 5 | 5.28 | 1.3650 | 0.26 | 216.45 | 3.5 | 183.7 |
| 6 | 2.46 | 0.6080 | 0.25 | 122.18 | 4.0 | 98.0 |
| 7 | 4.36 | 1.3340 | 0.31 | 167.99 | 3.8 | 164.6 |

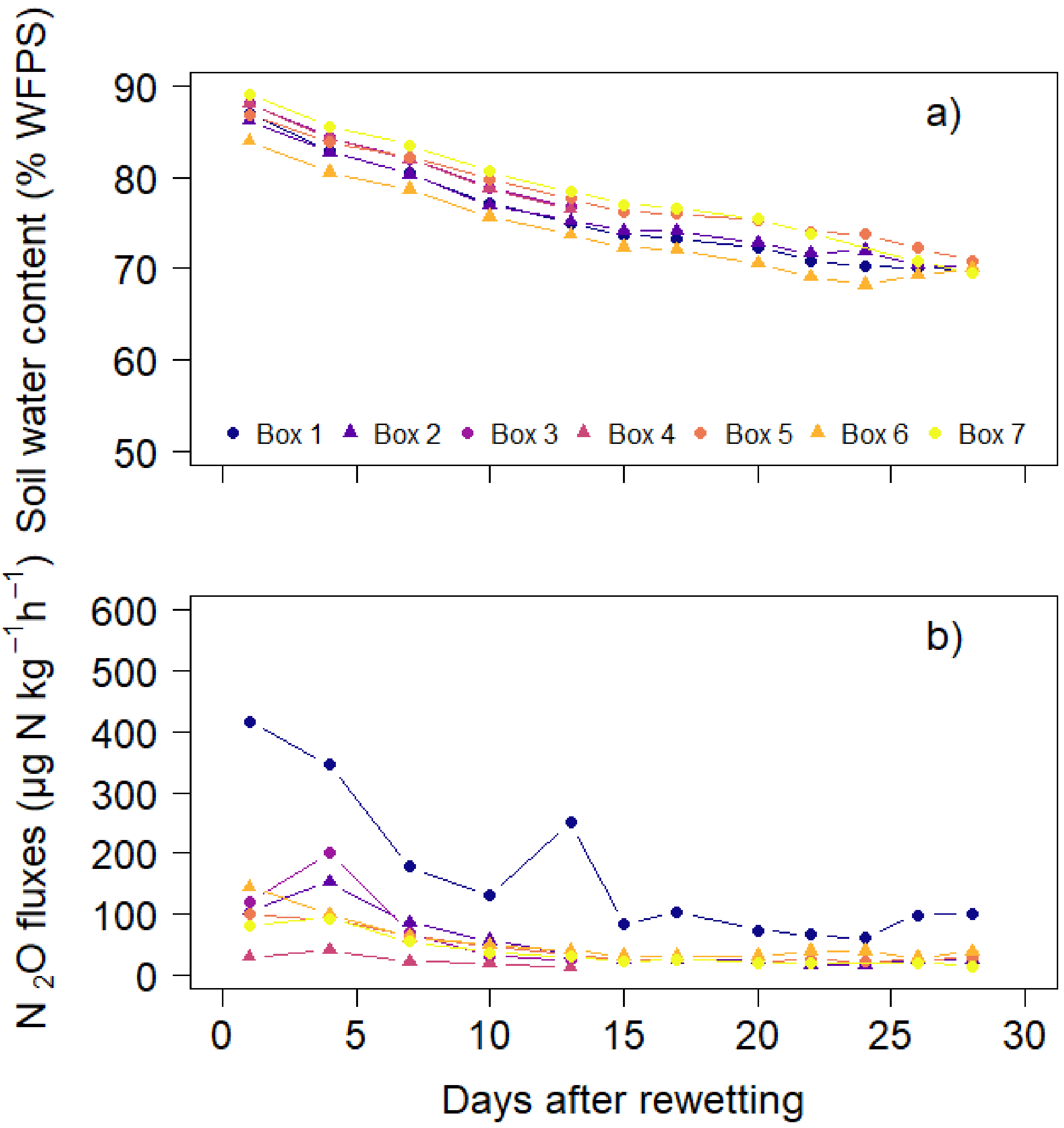


**Figure S1:** (a) soil water content (% water-filled pore space), and (b) $N_2O$ fluxes (µg $N_2O$-N $kg^{-1}$ $h^{-1}$) for each individual rhizobox before plant emergence.

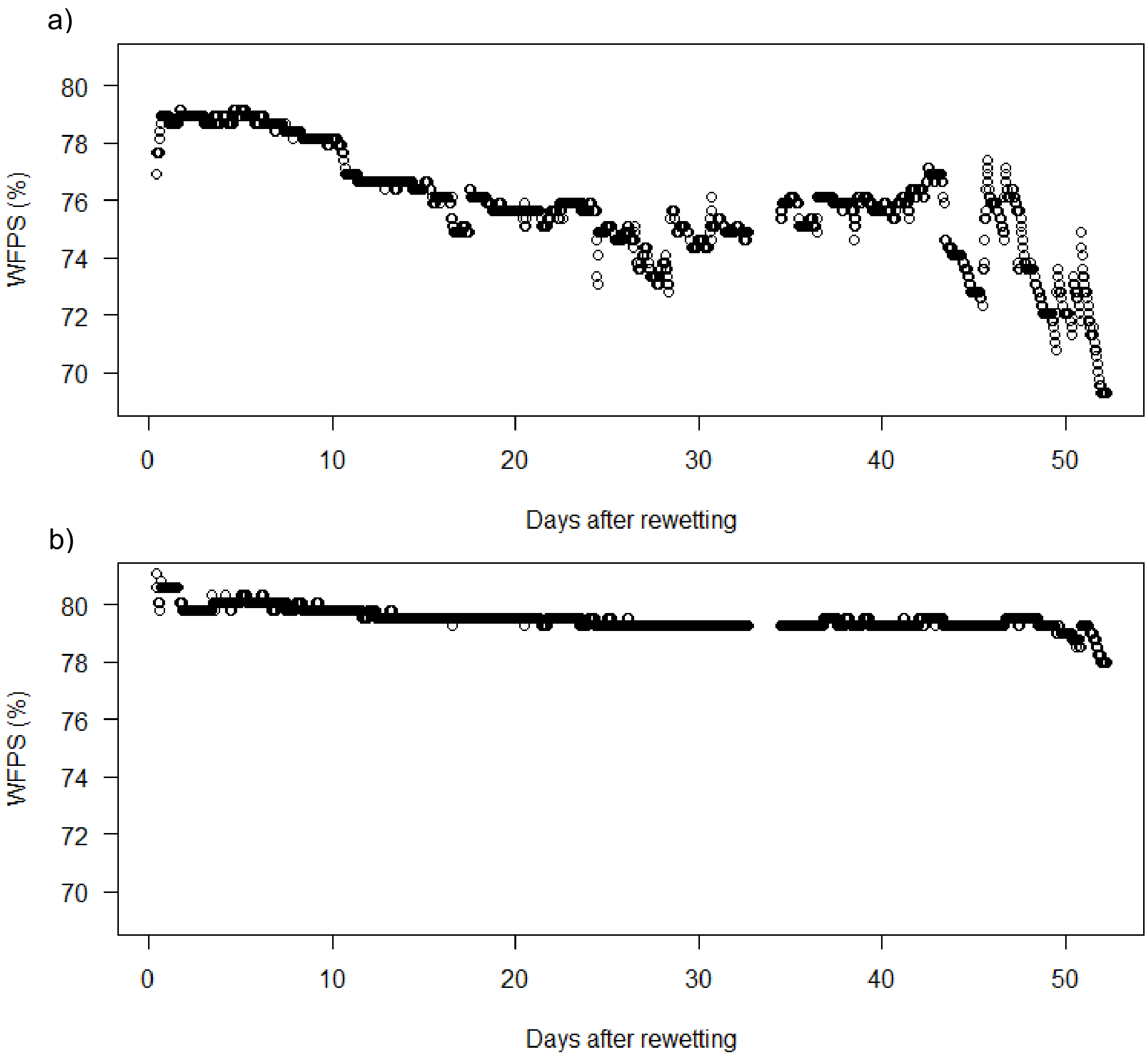


**Figure S2:** Soil moisture (in % WFPS) measured with FDR sensors in Rhizobox 7 at a) 5-10 cm and b) 30-35 cm depth.

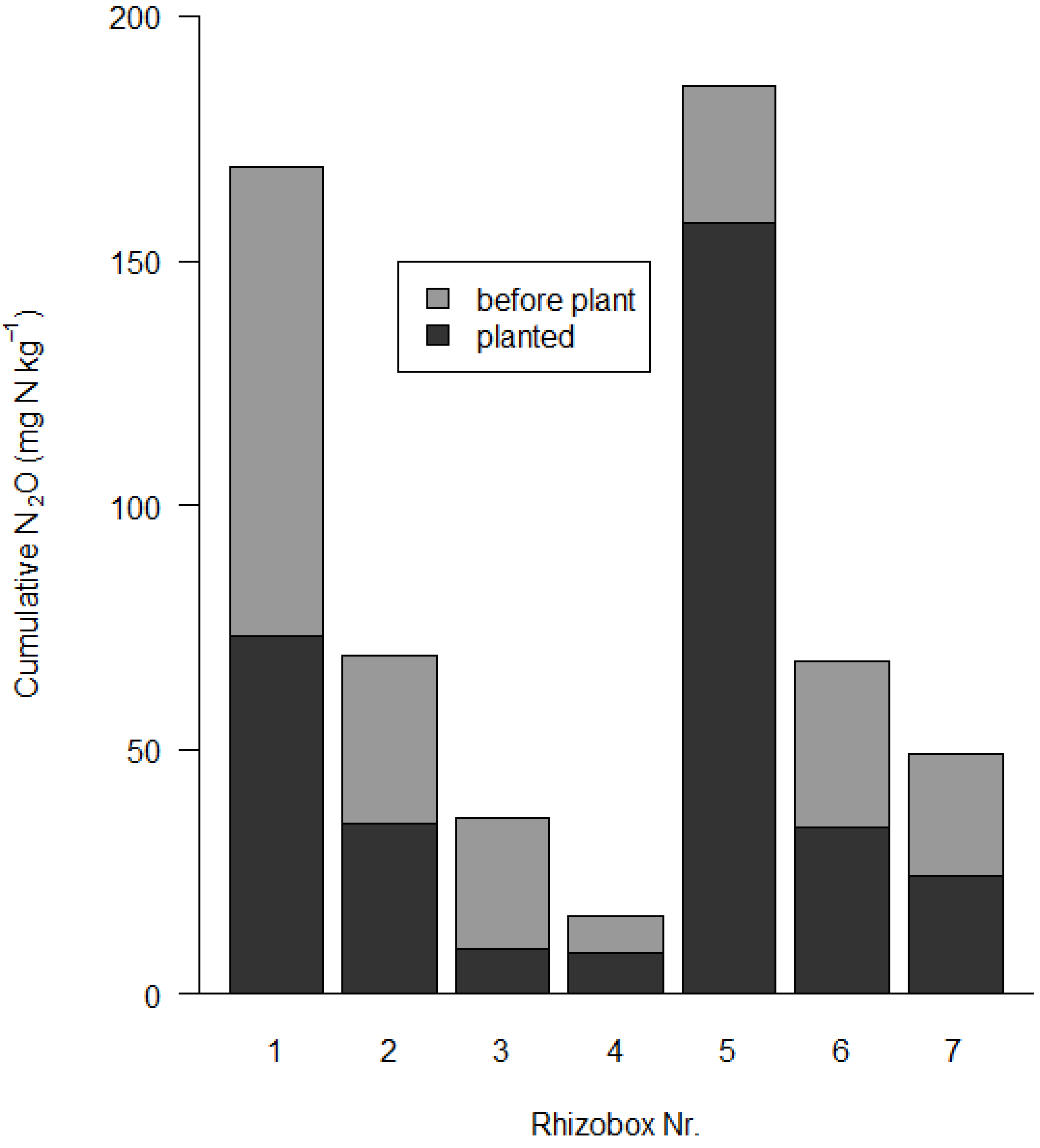


**Figure S3:** Cumulative $N_2O$ emissions of all rhizoboxes before and after plant emergence.

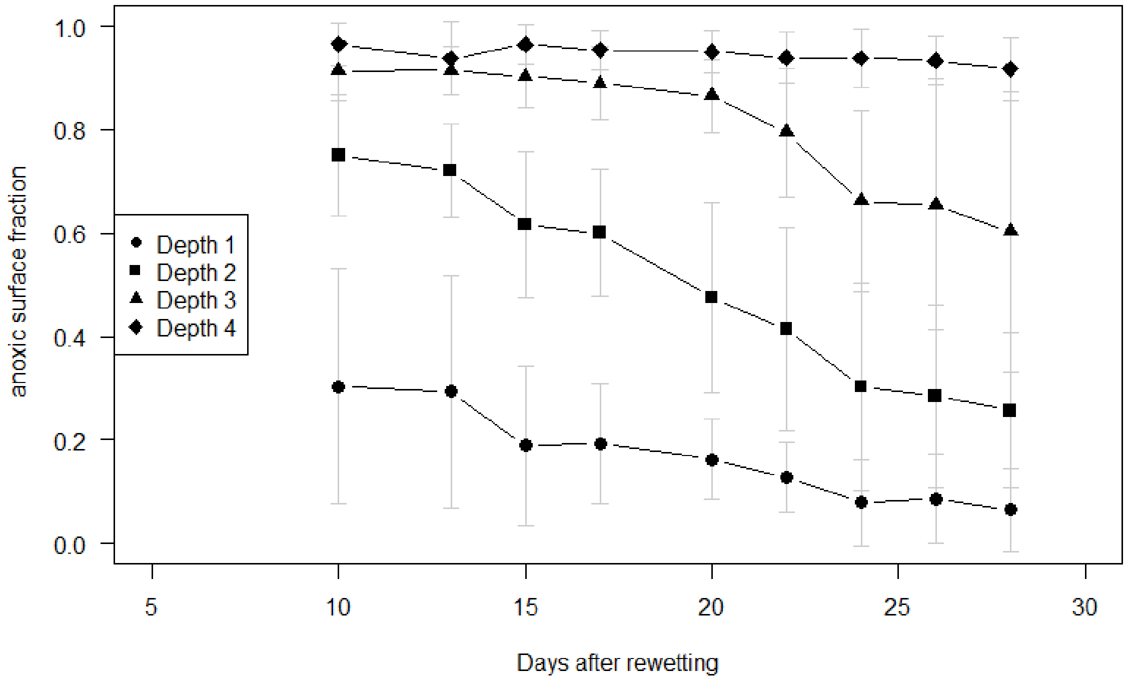


**Figure S4:** Fraction of anoxic area (< 10% $O_2$ air saturation = 2.1 % $O_2$) before plant emergence. Different symbols represent the four depths in the rhizobox. Mean ± standard deviation for n=7.

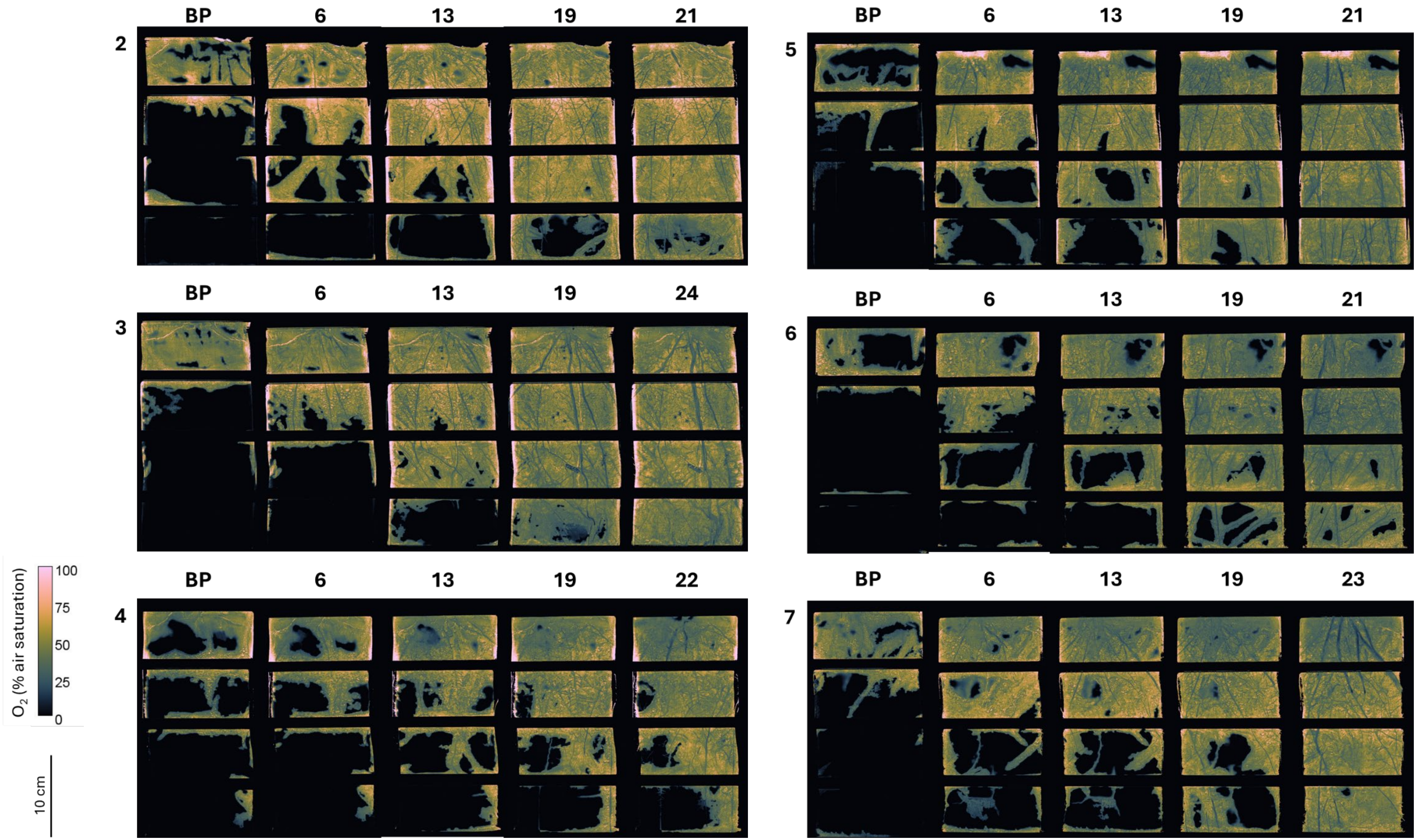


**Figure S5:** False color images of the $O_2$ concentrations measured with planar $O_2$ optodes. Numbers to the top left indicate the number of the rhizobox (i.e. the replicate). The numbers above the image indicate the day after plant emergence, BP = before plant emergence.

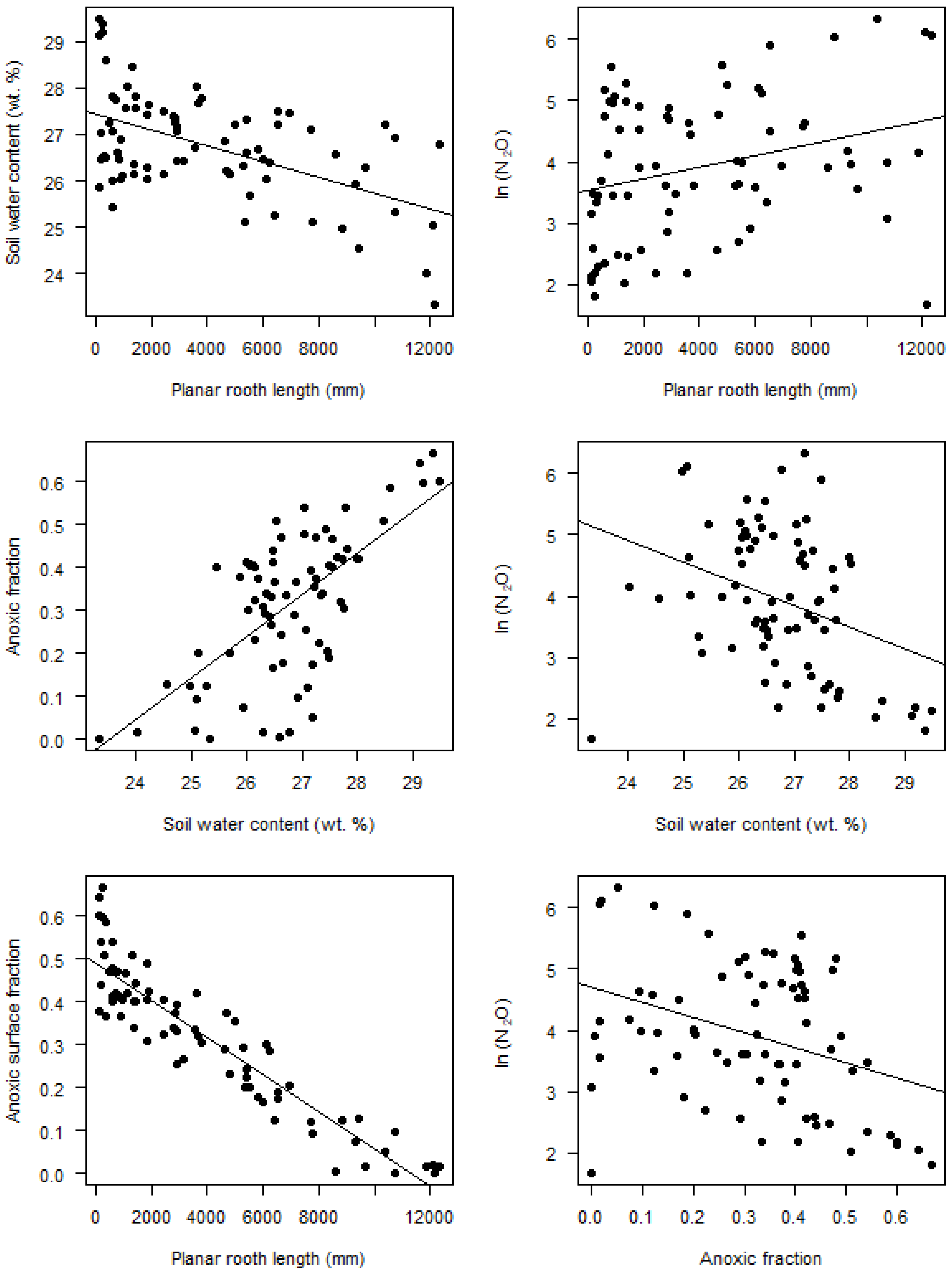


**Figure S6:** relationships between soil water content, planar root length, and anoxic fraction and between ln($N_2O$) and soil water content, planar root length, and anoxic fraction.

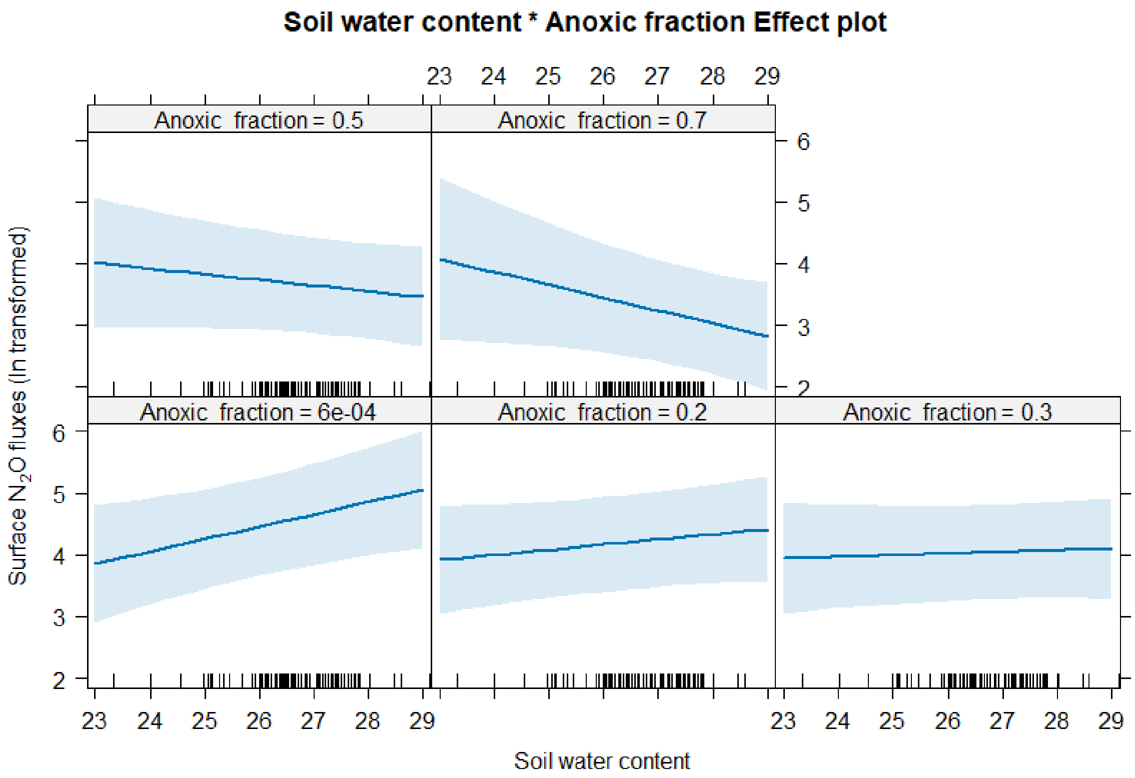


**Figure S7:** Interaction plots between surface $N_2O$ fluxes (ln transformed), gravimetric soil water content and anoxic fraction.

**Table S3:** Results of regression analyses (coefficients of determination (Adjusted $R^2$), *p*-values, and sample size *n*) of the relationship between soil water content, root length, anoxic surface fraction, and $N_2O$ surface fluxes (ln transformed) for the whole experimental phase

| Model | *n* | $R^2$ | *p*-value |
|---|---|---|---|
| *Simple linear regressions* | | | |
| Anoxic fraction ~ Soil water content | 124 | 0.635 | $< 2.2 \times 10^{-16}$ |
| ln($N_2O$) ~ Soil water content | 124 | 0.08944 | 0.0004349 |
| ln($N_2O$) ~ Anoxic fraction | 124 | 0.09642 | 0.0002639 |
| *Linear mixed effect model* | | | |
| ln($N_2O$) ~ Soil water content * Root length<br>Random effect: Rhizobox Nr (replicate)<br>Period: full experiment | 124 | 0.729 (cond.)*<br>0.151 (marg.)* | 0.0034 |

*For the linear mixed effect model (lme), the conditional $R^2$ is the variance explained by the entire model, including both fixed and random effects, while the marginal $R^2$ represents only the variance explained by the fixed effects

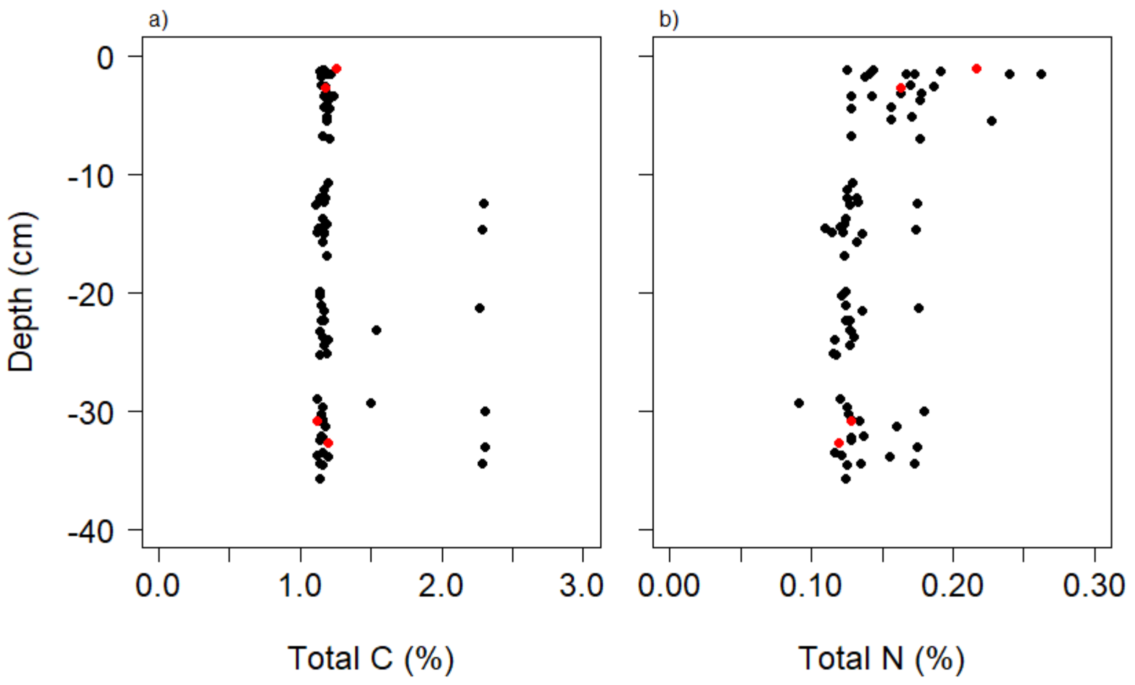


**Figure S8:** Depth distribution of (a) total soil C and (b) total soil N content in ROI. Each black point represents one ROI. Red circles represent ROI Nr. 1, 6, 45, and 77.

**Table S4:** Results of additional simple linear regression analyses (coefficients of determination (Adjusted $R^2$), p-values, and sample size *n*) of the relationship between N concentrations and abundance of nitrogen cycling genes in ROI.

| Model | *n* | $R^2$ | *p*-value |
|---|---|---|---|
| **16S ~ $NH_4^+$** | **77** | **0.3235** | **$4.088 \times 10^{-08}$** |
| ***nirK* ~ Depth** | **77** | **0.3629** | **$4.078 \times 10^{-09}$** |
| *nirS* ~ Depth | 77 | 0.005957 | 0.2314 |
| ***nosZI* ~ Depth** | **77** | **0.3449** | **$1.188 \times 10^{-08}$** |
| ***nosZII* ~ Depth** | **77** | **0.5349** | **$2.557 \times 10^{-14}$** |

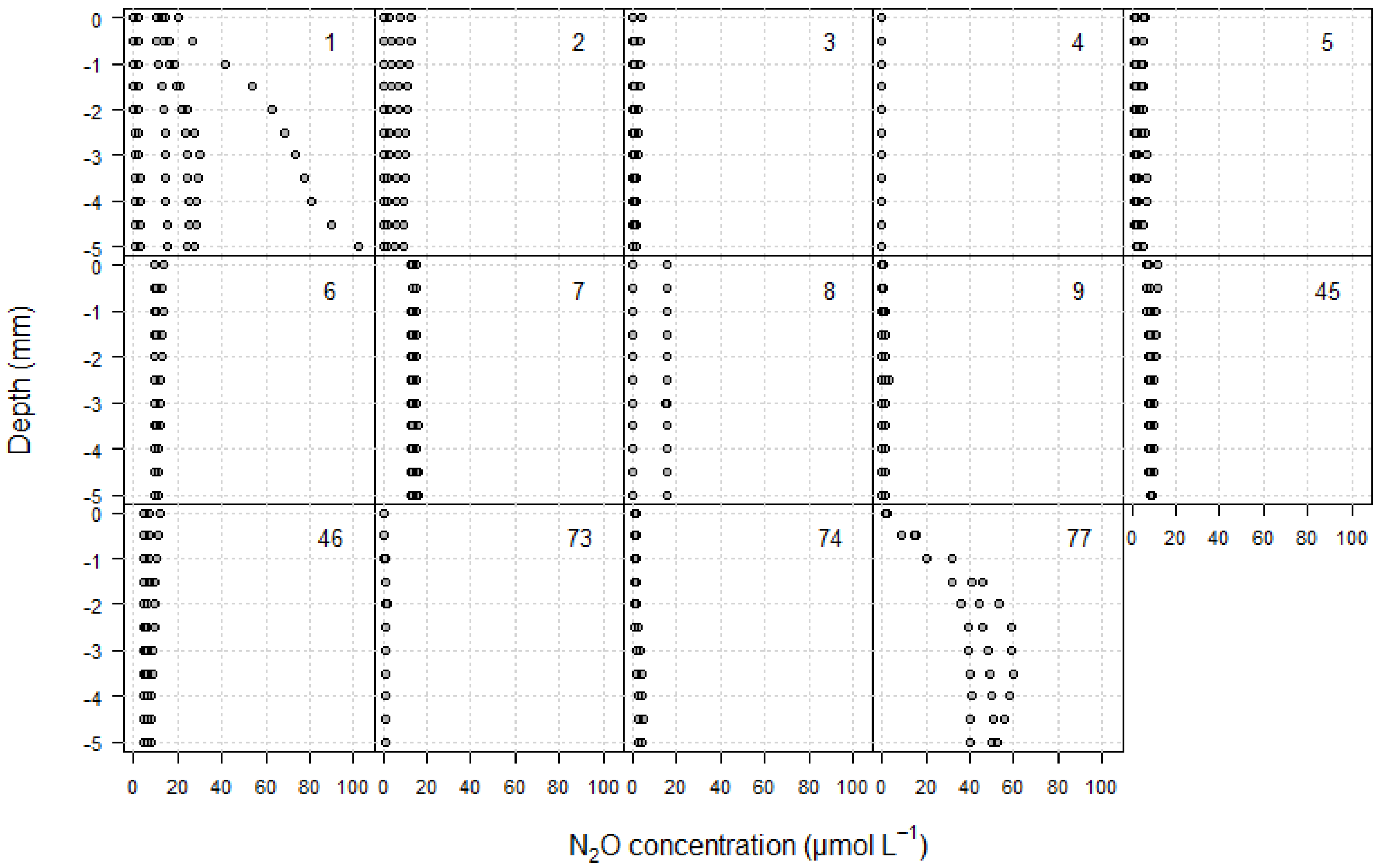


**Figure S9:** $N_2O$ concentration depth profiles measured with $N_2O$ microsensors at the end of the experimental period. The number in the upper right corner of each panel refers to the number of the ROI in which the measurements were done.

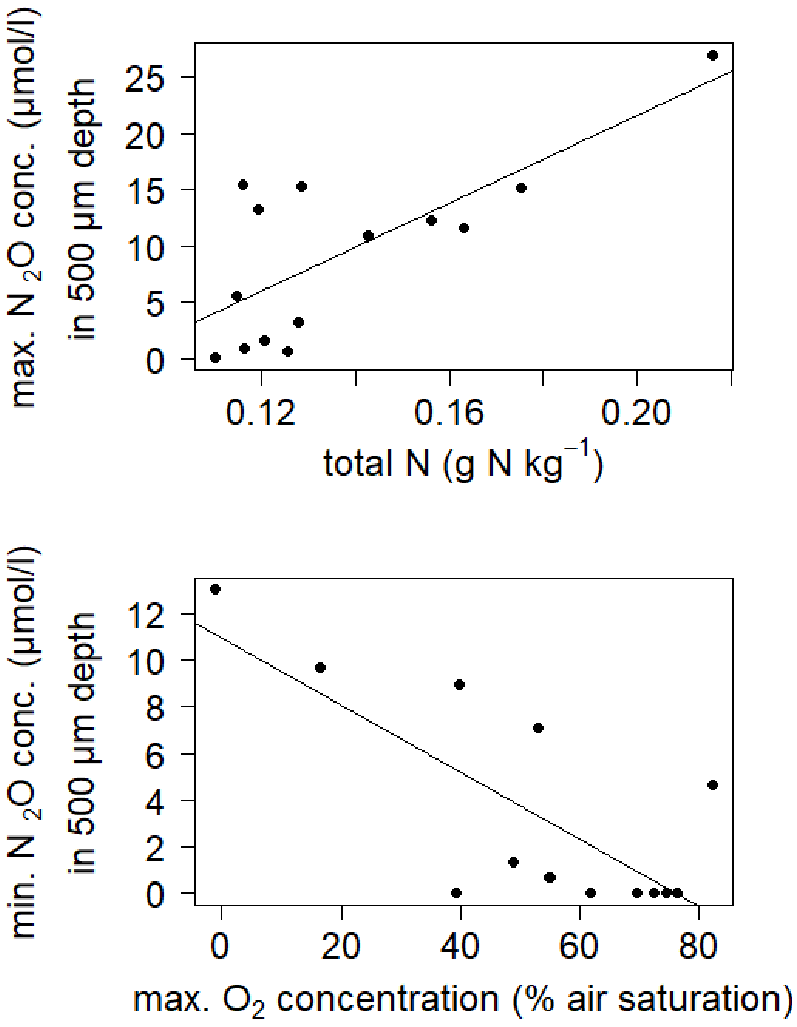


**Figure S10:** linear regressions between $N_2O$ concentrations measured with microsensors in 500 µm depth and total N or maximum $O_2$ concentration in ROI.
